# Spatial structure could explain the maintenance of alternative reproductive tactics in tree cricket males

**DOI:** 10.1101/2024.03.25.586515

**Authors:** Mohammed Aamir Sadiq, Ananda Shikhara Bhat, Vishwesha Guttal, Rohini Balakrishnan

**Affiliations:** Centre for Ecological Sciences, Indian Institute of Science, Bangalore, India; Indian Institute of Science Education and Research, Pune, India; Institute of Organismic and Molecular Evolution (iomE), Johannes Gutenberg University, 55128 Mainz, Germany; Institute for Quantitative and Computational Biosciences (IQCB), Johannes Gutenberg University, 55128 Mainz, Germany

**Keywords:** alternative reproductive tactics, individual-based model, fitness, spatial structure, frequency dependence, population density

## Abstract

Trait polymorphisms are widespread in nature, and explaining their stable co-existence is a central problem in ecology and evolution. Alternative reproductive tactics, in which individuals of one or more sex exhibit discrete, discontinuous traits in response to reproductive competition, represent a special case of trait polymorphism in which the traits are often complex, behavioural, and dynamic. Thus, studying how alternative reproductive tactics are maintained may provide general insights into how complex trait polymorphisms are maintained in populations. We construct a spatially explicit individual-based model inspired from extensively collected empirical data to address the mechanisms behind the co-existence of three behavioural alternative reproductive tactics in males of a tree cricket (*Oecanthus henryi*). Our results show that the co-existence of these tactics over ecological time scales is facilitated by the spatial structure of the landscape they inhabit, which serves to equalize the otherwise unequal mating benefits of the three tactics. We also show that this co-existence is unlikely if spatial aspects of the system are not considered. Our findings highlight the importance of spatial dynamics in understanding ecological and evolutionary processes and underscore the power of integrative approaches that combine models inspired from empirical data.

## Introduction

The persistence of polymorphisms in populations has been a central theme in evolutionary biology. Alternative reproductive tactics (ARTs) are a class of polymorphisms that evolve in response to competition for mates and manifest as discrete morphological, physiological, and/or behavioural traits that can be expressed by individuals of either sex (Brockmann & Taborsky, 2008). For example, male guppies (poecilid fish) are known to either court or forcefully copulate with females (Bisazza & Pilastro, 1997). Likewise, female digger wasps (*Sphex ichneumoneus*) either build their nests to lay eggs or parasitise other females’ nests (Brockmann, et al., 1979). Male field crickets either call to attract females or behave as satellites (Cade, 1981).

Traditionally, the persistence of ARTs has been understood with reference to their mode of expression. ‘Genetically determined’ ARTs are entirely controlled by a few loci and are thought to be maintained in populations through mechanisms such as neutrality (Shuster & Wade, 1991), spatiotemporally varying selection pressures (Mazer & Damuth, 2001), negative frequency-dependent selection (Mart R Gross, 1996) and heterozygote dominance (Krüger et al., 2001). On the other hand, ‘condition-dependent’ ARTs are conditionally expressed depending on an organism’s intrinsic state or extrinsic environment and can persist as long as different tactics are optimal in different conditions and all these different conditions frequently occur in nature (Brockmann & Taborsky, 2008). However, the classification of ARTs into genetic and conditional is controversial, as it is increasingly clear that their phenotypic expression is regulated by a combination of genetic and environmental (intrinsic or extrinsic) factors (Brockmann, 2001). Understanding how the demographic and physical environment affect the fitness of ARTs independent of their mode of expression may yield more insights into coexistence of ARTs and contribute to a general theoretical framework for their persistence.

Trait frequency and population density are important demographic factors that affect the fitness of alternative tactics and may determine whether a polymorphism can be maintained in a population (Gross, 1996; Kokko & Rankin, 2006). Negative frequency-dependent selection, wherein the fitness of an ART decreases as its frequency of expression in the population increases, is thought to be a common mechanism for the coexistence of ARTs (and polymorphisms in general), especially in evolutionary game theory (Wolf & Waltz, 1993). This phenomenon can lead to the maintenance of ARTs at a constant (Gross, 1991; Wolf & Waltz, 1993) or oscillating frequency (Takahashi & Kawata, 2013) based on system specifics.

Population density can also affect the fitness of ARTs and determine which tactic (if any) has the highest fitness in the population. For example, in *Sancassania berlesei,* a species of acarid mites, males can either fight or scramble to gain mates. Artificial selection experiments show that the fitness of fighter morphs in this species decreases with an increase in population density (Michalczyk et al., 2018), possibly due to a higher propensity for harmful interactions with conspecifics. In field populations of the European field cricket (*Gryllus campestris*), silent males are more successful at mate-finding than calling males at high population densities, with the reverse holding true at low population densities (Hissmann, 1990) due to a change in encounter probabilities. Furthermore, both theoretical (Eadie & Fryxell, 1992) and empirical (Takahashi & Kawata, 2013) studies have shown that population density can have an amplifying effect on the strength of frequency-dependent selection and thus facilitate coexistence.

Spatial structure in the environment can also impact the fitness of alternative tactics and change the potential for the maintenance of polymorphisms. For example, fighter males of the mite *Rhizoglyphus echinopus* show lower fitness in complex 3D habitats, possibly due to the limited movement ability of the fighter males and the spatial distribution of females (Tomkins et al., 2011). Heterogeneity brought about by a spatially structured landscape provides an opportunity for multiple phenotypes to coexist via local adaptation (Gray et al., 2008; Korona et al., 1994; Stein et al., 2014; Zahnd et al., 2021). Computational and analytical models in the context of migration and cooperation show that movement and spatial structure can promote trait polymorphism and even oscillatory states (Guttal & Couzin, 2010; Joshi et al., 2017). In spatial models of competitive Lotka-Volterra communities, incorporating ‘steric’ structures that obstruct dispersal within the landscape can lead to more localized interspecific interactions in space, which in turn can facilitate ecological coexistence (Ursell, 2021). In nature, habitats are rarely isotropic and steric obstructions to movement abound, meaning that incorporating such heterogeneity is vital to making inferences about real-life populations in the field.

The reproductive life-history of sexual organisms often involves signal-receiver dynamics where individuals emit a signal to improve detectability or attractiveness to members of the opposite sex. In organisms that show acoustic signal-receiver dynamics, males can adopt multiple ARTs such as calling and satellite behaviours. Steric structures in the habitat can change the effective distance up to which a signal can be transmitted (Endler, 1992; Jain & Balakrishnan, 2012). Steric structures can also serve as microhabitats and affect the spatial distribution and movement patterns of signallers and receivers. This is especially relevant for species which exhibit ARTs: different tactics may employ diverse signaling and movement behaviors, and steric structures could differentially affect the fitness of different ARTs, with potential implications for the maintenance of polymorphisms over evolutionary time. The effect of spatial structure on the coexistence of such behavioural ARTs has, however, received relatively little attention.

In the tree cricket *Oecanthus henryi*, as in many cricket species, males call by stridulating with their forewings to attract mates (Walker, 1957). Alternatively, males can choose to remain silent (Torsekar & Balakrishnan, 2020). A third ART (‘baffling’) exists in which males construct an acoustic baffle (amplifier) using a leaf from their host plant (*Hyptis suaveolens*) and call through this structure (Deb et al. 2020). Baffling amplifies the male call, increasing the distance over which the male can be perceived by phonotactic females (Prozesky-Schulze et al., 1975) and increasing the attractiveness of a male to phonotactic females (Deb et al., 2020). Given these obvious mating advantages, one may expect that all males should baffle. Despite this, only a fraction of males are bafflers in natural populations of *O. henryi* (Deb, 2017). We explore this conundrum using an IbM based on empirical data. Specifically, we investigate whether mechanisms such as negative-frequency dependence, population density and spatial structure could i) prevent baffling from always being advantageous over other tactics, and ii) facilitate the coexistence of baffling, calling (without baffling), and silent mating tactics in *O. henryi*.

Individual-based models (IbMs) are a powerful tool to study the effect of spatial structure as they explicitly model the local interactions that provide a mechanistic basis for observable patterns at the population level. IbMs further allow us to capture emergent properties resulting from non-linear interactions which may not be feasible using analytical models. IbMs of calling and satellite cricket males show that their fitness is likely strongly affected by population density but that frequency-dependence plays a relatively minor role (Rowell & Cade, 1993; Walker & Cade, 2003). Similar models have also shown that both calling and satellite tactics can coexist despite high levels of parasitism by an acoustically orienting parasitoid (Rotenberry & Zuk, 2016). These studies do not, however, incorporate the spatial structure of the habitat in their simulations.

## Methods

We use a spatially explicit, individual-based model of *Oecanthus henryi* in which we specify sex and tactic-specific rules for each individual. The values of the parameters used in the model are primarily inspired and abstracted from the observations made on *O. henryi*. Description of model parameters, their values as well as their empirical sources are stated in the following sections and summarized in Table 1. For instances where we used parameterized distributions, the goodness of fit of distributions to the empirical data are summarized in Table 2. Here, we summarise the broad outline of the model and the details are given in the sub-headings that follow.

**Table 1.**
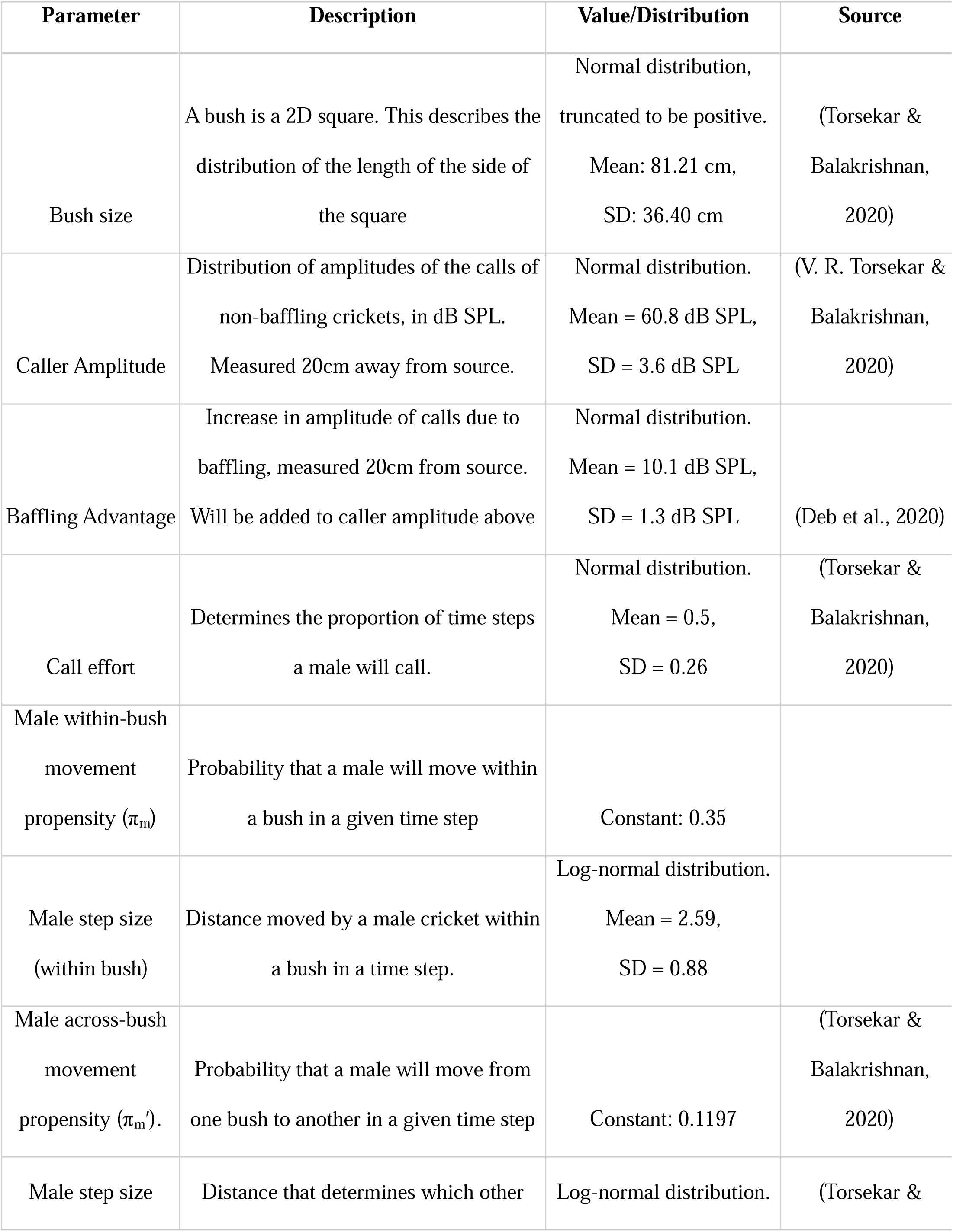
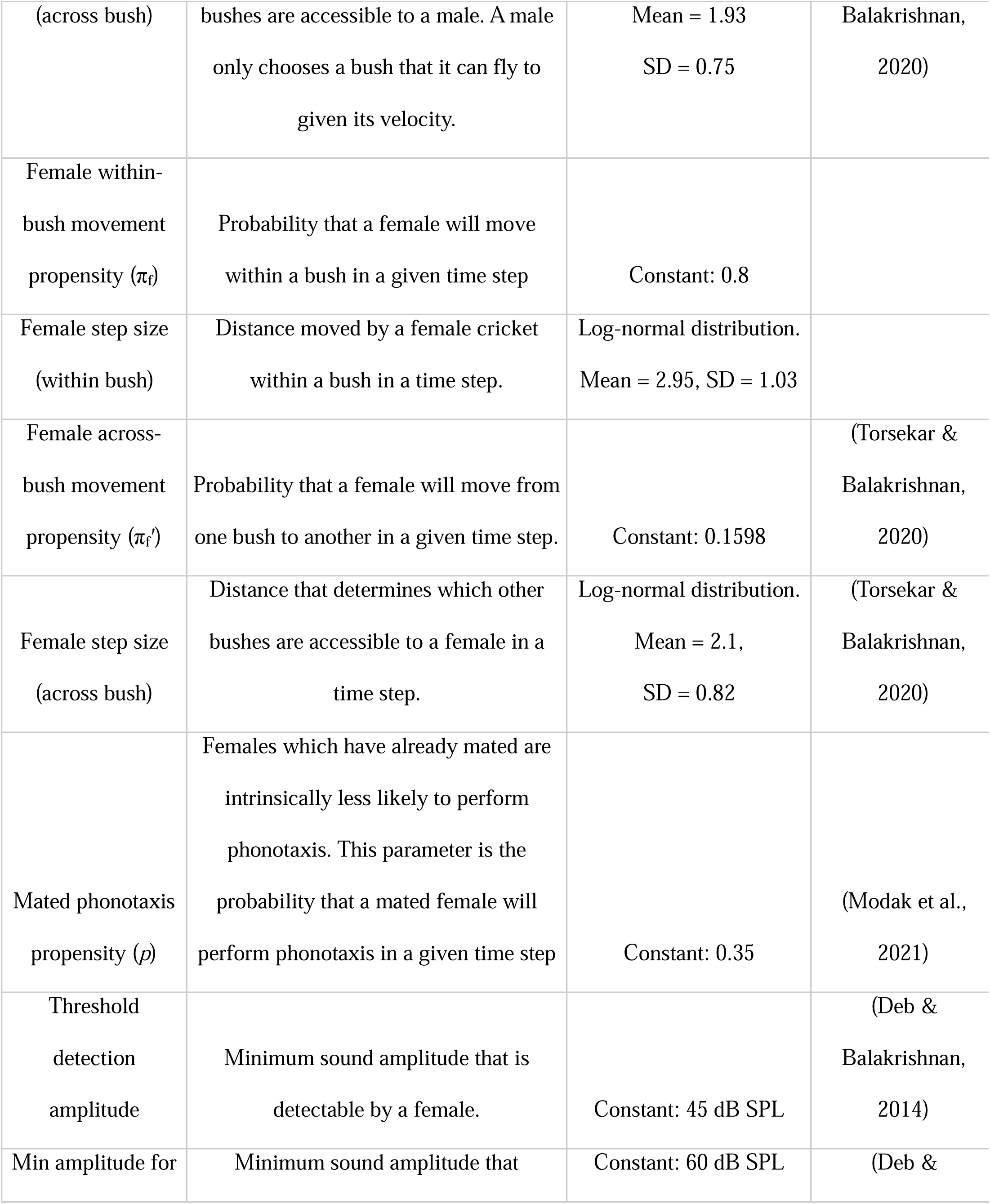
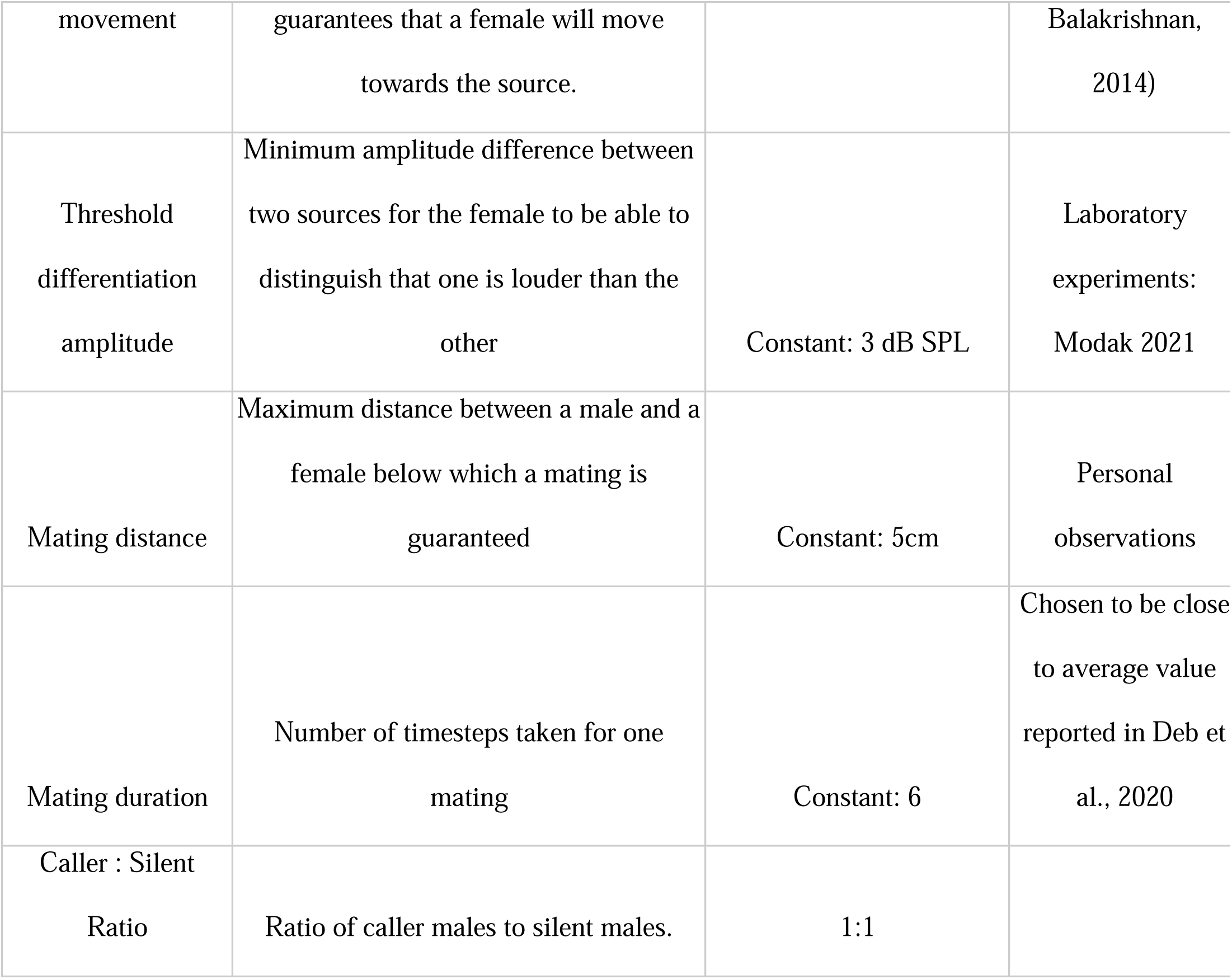
List of all static model parameters and their empirical sources.

**Table 2.** Summary of parameter distributions for male and female movement, male calling effort and bush widths (i.e., bush sizes). If the observed Kolmogorov-Smirnov (KS) distances are less than the KS distance for α=0.05, then the chosen theoretical distributions are similar to the empirical distributions.

| **Variable** | **Chosen theoretical distribution** | **KS distance for $\alpha=0.05$** | **Observed KS distance** |
| --- | --- | --- | --- |
| Male step size across bush | *Lognormal(1.9,0.7)* | 0.14 | 0.08 |
| Female step size across bush | *Lognormal(2.1, 0.8)* | 0.11 | 0.07 |
| Male step size within bush | *Lognormal(2.6,0.9)* | 0.15 | 0.07 |
| Female step size within bush | *Lognormal(2.9,1.0)* | 0.124 | 0.111 |
| Male calling effort | *Normal (0.5, 0.26)* | 0.06 | 0.08 |
| Bush width | *Normal(81.2, 36.4)* | 0.09 | 0.04 |

### Model Overview

Our model consists of individual crickets positioned in a 2-D square arena with reflective boundaries. Each simulation consists of 500 individuals (1:1 sex ratio) and is run for 72 time-steps, where each time step corresponds to 10 minutes of real time and the total simulation time captures one night (12 hours). Each male is assigned one of three tactics (silent, calling, baffling) and this tactic is fixed for the entire duration of a simulation. The males and females move and interact according to rules that reflect empirical findings (described later). We vary the proportion of bafflers and the spatial structure. We also vary population density (*d*) by varying the size of the square arena. We then evaluate the average mating success of each tactic employed by individual males. The details of the model are described below.

### Spatial Structure

Individuals inhabit a large two-dimensional, square arena, whose size is determined according to the specified population density *d* (no. of individuals per square metre). We consider a range of densities from 0.05 to 1 individual per sq m (i.e., *d* = { 0.05, 0.25, 0.5, 0.75, 1 }). The range of population densities chosen in our model was inspired by inter-individual distances measured in the field ((Deb & Balakrishnan, 2014). Tree crickets are generally found on trees or bushes, and the species we study (*O. henryi*) is found mainly on *Hyptis suaveolens* bushes. To check for the effect of spatial structure in the form of bushes, we modeled two distinct kinds of habitats (as illustrated in Fig. 1).

**Fig. 1.**
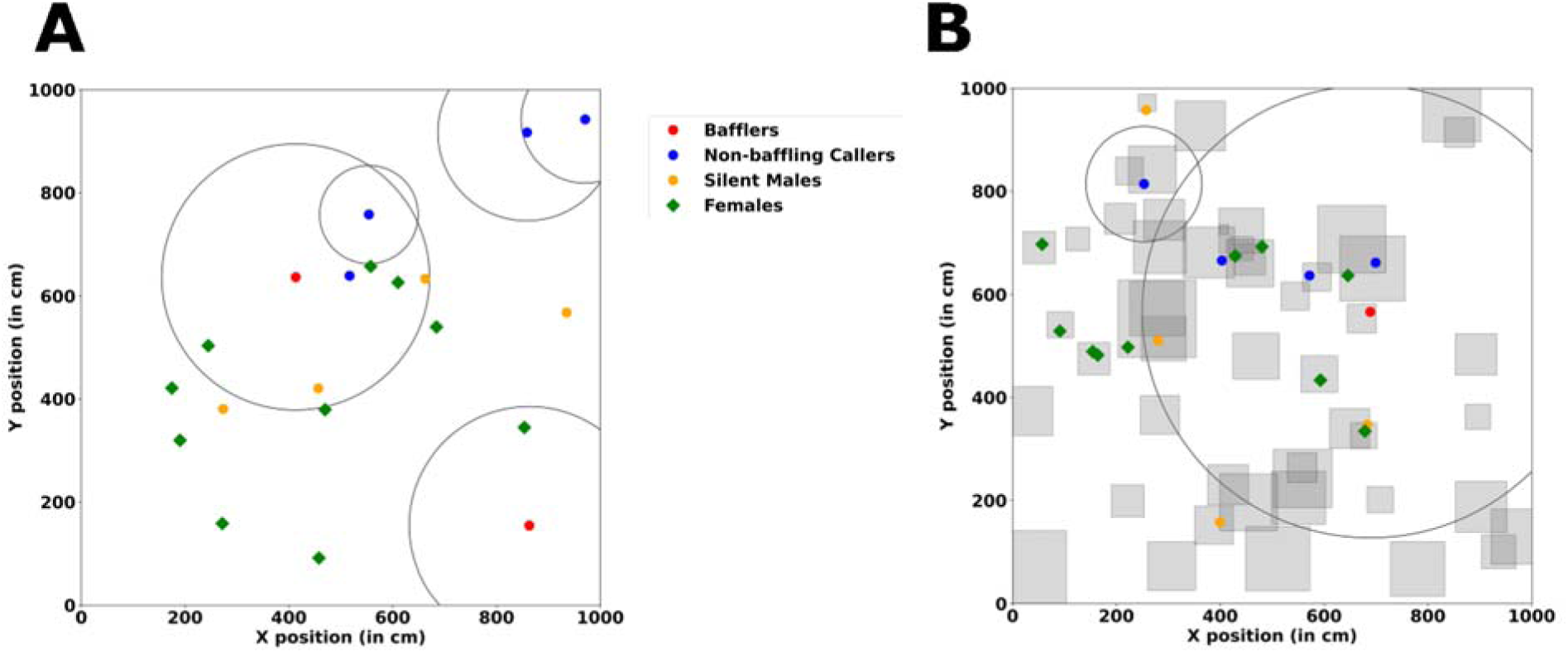
A representative illustration of the simulation arena. A) A homogeneous simulation arena. B) A heterogeneous simulation arena with squares indicating bushes. The circles indicate the radial distance at which sound pressure level (SPL) of a male caller/baffler (coloured dots at the centre of circles) drops to 45 dB (hearing threshold of females).

### Homogeneous habitat

As a model for unstructured habitats, we construct a homogeneous habitat where we do not account for the presence of bushes and the individuals are free to move throughout the landscape (Fig. 1A). To initiate the simulation, individuals are positioned throughout the landscape according to a random uniform distribution. The homogeneous habitat acts as a null model to compare with the effects of structured habitats.

### Structured habitat

As an approximation of the ecological reality of the habitat we incorporate bushes in our landscape. Bushes are abstracted as closed squares within the landscape (Fig. 1B). Individuals can either move within the same bush or instantaneously move from one bush to another, but cannot exist outside of a bush at any given time step. The presence of bushes therefore brings about heterogeneity in the landscape. The position of the centres of the bushes was determined by a uniform random distribution whose limits were determined by the edges of the landscape. The number of bushes is determined by the spatial density of bushes (no. of bushes per sq. m), denoted by ρ. In structured habitats, the expected number of animals on a bush is given by d/ρ where d is population density and ρ is bush density. Thus for a bush density value of 1, the number of individuals on a bush can range from 0.05 to 1 for population densities ranging from 0.05 to 1.We tested for the influence of the spatial density of bushes by varying ρ from 0.5 to 2 bushes per sq. m. In the field, bushes are present at a mean density of about 1.6 bushes per square meter but this number is subject to substantial seasonal variation (Torsekar et al. 2019).

### Acoustic properties of the landscape

Since male tree crickets emit acoustic signals to attract mates, the attenuation properties of sound within the landscape are important for their reproductive ecology. In our model, we assume that the sound pressure level decays according to the laws of spherical spreading. Our assumption of spherical spreading of sound reflects the fact that the presence of bushes in the field causes negligible excess attenuation (Deb & Balakrishnan, 2014). The intensity of sound is measured in terms of sound pressure level (SPL). According to the laws of spherical spreading, *i.e.,* the transmission loss (*L*) of a signal in dB SPL at a distance *d* metres away from the source is assumed to be given by:

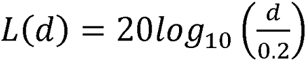

The constant 0.2 reflects the fact that the empirical measurements of sound amplitudes that we used to parameterize the simulation are measured from a distance of 0.2 m (20 cm) away from the source. The reflective boundary conditions on animal movement ensure that individuals on one end of the landscape do not listen to individuals signaling from the opposite end, thus preventing possible boundary effects due to unphysical acoustic transmission.

### Individual attributes

Males can perform one of three tactics, and the tactic assigned to them is fixed for the duration of the simulation run:

1. **Bafflers** can call but their location is fixed for the duration of the simulation.
2. **Callers** can both call and move randomly through the landscape.
3. **Silent males** execute random movements but do not emit any calls.

To check for the effect of proportion of bafflers on the mating success of the ARTs, we varied the proportion of bafflers in the population. The remaining males are equally partitioned into the caller and silent ARTs, in accordance with empirical findings which show that a male is equally likely to call or remain silent on a night (Sadiq et al., 2023, submitted).

Since the ARTs differ with respect to both signaling and movement, we elucidate them separately.

### Signaling

Silent males do not emit any calls. Callers and bafflers were modeled as point emitters of spherically symmetric sound. They have an associated call effort, defined as the proportion of time steps (out of 72) for which they emit a call. Based on empirical work (Torsekar & Balakrishnan, 2020), the call effort of each signaling male is drawn from a truncated normal distribution with a mean 0.5 and SD 0.26. Since call effort must be a number within the interval (0,1], we resampled if the drawn call effort was outside this interval. Once the call effort is determined for each individual, the specific time steps in which a male calls are randomly assigned. In addition, we assume that a male will not call when it is mating. Each signaling male also has an associated amplitude for its call. The SPL of a caller is drawn from an *N(60.8 dB, 3.6 dB)* distribution following empirical data (Deb et al., 2020). If the male is a baffler, it receives an additional amplitude boost due to its baffle relative to the non-baffling caller. The excess amplitude due to the baffle is also drawn from an *N(10.1, 1.3)* distribution (Deb et al., 2020).

### Movement

In our simulations, bafflers are not allowed to move since baffling requires males to make a hole and call from a leaf. Silent males and callers can move, but we assume that callers do not move while emitting a call. The extent and likelihood of movement differ based on whether the movement is within a bush or across bushes. Males can perform only one of the two movements (within or across bush movement) in a time step. The likelihood of movement within a bush is denoted by π_m_. In this case, the distance (in cm) a male moves is drawn from an *LN(2.59, 0.88)* distribution, i.e., a log-normal distribution with mean 2.59 and standard deviation 0.88. The parameters for the log-normal distribution are derived from empirical data (Sadiq et al. 2023, submitted). If a male does not move within a bush, it may instead move across bushes with probability π_m_^′^. Therefore, the net likelihood of across-bush movement is given by [1-π_m_] π_m_′ (i.e., probability of not moving within a bush multiplied by the probability of moving across bushes) and is lower than the likelihood of movement within a bush. Our assumption of high within-bush-movement and low-across-bush-movement is motivated by the high site fidelity previously observed in males of *O. henryi* (Deb & Balakrishnan, 2014) within nights and is common to other taxa as well (Kleyla & Dodson, 1978; Pittman et al., 2008). For movement within the bush, the direction is chosen randomly. When individuals execute movement across bushes, they move to a randomly chosen bush whose center is within a distance λ_m_ from the centre of the bush the male currently occupies. λ_m_ is drawn from a *LN(1.93, 0.75)* distribution each time the male moves across bushes (Torsekar & Balakrishnan, 2020). The males assume a random position within this new bush. In homogenous habitats, males move according to within-bush parameters. While it is interesting to assume that silent males behave as satellites and show directed movement towards calling males, we assumed randomness in the directionality of movement for both calling and silent males for the sake of simplicity.”

### Females

Females are silent and perform two types of movement: 1) directed movement in response to male signals, termed as phonotaxis; 2) movement in a random direction when not within the sound field of calling males. Laboratory experiments indicate that mating status (virgin or mated) can affect female movement (Modak et al., 2021). In line with previous studies (Cade, 1993), we therefore implemented different relative probabilities of phonotaxis and random movement for females in our simulations based on their mating status.

### Movement

Females show a high likelihood of movement within a bush, and we denote this likelihood by π_f_. In this case, the distance (in cm) moved by the female in a timestep is drawn from a *LN(2.95, 1.03)* distribution. If a female does not move within a bush, it may instead move across bushes with probability π_f_′. Therefore, the net likelihood of across-bush movement is given by [1-π_f_]π ′, i.e., probability of not moving within a bush multiplied by probability of moving across bushes and is lower than the likelihood of movement within a bush. When individuals execute movement across bushes, they move to a bush whose center is within a distance λ_f_ from the centre of the bush the female currently occupies. λ_f_ is drawn from a *LN(2.1, 0.82)* distribution each time the female moves across bushes, in accordance with empirical data (Torsekar & Balakrishnan, 2020). The females assume a random position within this new bush. In homogenous habitats, females move according to within-bush parameters. The direction of movement is random unless the females perform phonotaxis.

### Phonotaxis

At every time step, the probability that a female samples her acoustic landscape is dependent on her mating status and is given by a value of 1 for virgin females and a value of 0.35 for mated females (Modak et al., 2021). Once the female samples her landscape, the probability that a female responds to a signal of perceived intensity *I* is given by:

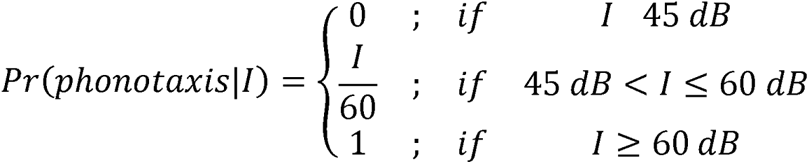

45 dB is the minimum SPL below which females are guaranteed to not respond phonotactically. 60 dB is the minimum threshold above which females are guaranteed to respond phonotactically. The implementation of this threshold response is supported by empirical data (Deb & Balakrishnan, 2014).

If a female is simultaneously exposed to two or more sounds whose perceived intensity is greater than 45 dB, it preferentially moves towards the loudest one. In line with laboratory experiments, females were additionally assumed to have an amplitude resolution of 3 dB meaning that they do not distinguish between two sounds whose perceived intensities differed by less than 3 dB (Modak, 2021). If exposed to two or more sounds whose perceived intensities are less than 3 dB apart, they choose one at random and move towards it.

Since females show high bush fidelity (see *Movement* section under *Females*), they first listen for sound sources within their own bush, and only listen for sounds from neighbouring bushes if there are no sufficiently loud sounds being emitted from their own bush. For calculating the perceived SPL of sound(s) for a listener positioned on a different bush, we sum the SPL of each individual present in the bush (after making the appropriate conversions from dB SPL to Pa for addition) to obtain the total intensity of sound emanating from the bush. Thus, we implicitly assume that males call in phase such that calls constructively interfere with each other. The perceived SPL is then calculated after accounting for the transmission loss due to the distance between the centres of the two bushes. Individuals from bushes that are greater than 10 m away from the focal bush are considered inaudible to the female since sounds from 10 m away are sub-threshold due to transmission loss.

### Mating

From empirical observations and experiments, we know that females do not reject males upon encounter, and typically mate with ∼ 90% probability (Modak et al. 2020). Consequently, we assumed in our model that two individuals of the opposite sex mate if they are within 5 cm (equivalent to ∼5 body lengths) of each other. If there are multiple males within 5 cm of a single female, she chooses a male randomly for mating. While mating, neither individual can move or engage in any other matings and males cannot emit calls. A mating event lasts for six time steps, which corresponds to a typical mating period (i.e., 1 hour in real-time). At the end of the mating period, we re-position the female to a random location within the landscape and assign it a ‘mated’ status.

The model was written in Python 3.9.7 (Van Rossum and Drake, 1995), and utilized the Python packages numpy 1.20.3 (Harris et al., 2020), scipy 1.7.1 (Virtanen et al., 2020), random 3.9.7, and multiprocessing 0.70.14. (McKerns et al., 2011).

### Quantitative inferences

We employed linear regressions to quantify the effect of proportion of bafflers, population density and spatial structure. The mating success of a tactic in a simulation run was measured as the average number of mates obtained by a male performing that tactic which implies that it is possible for males to have a mating success greater than 1. Since our simulations were stochastic, there was significant variation between individual runs of a simulation for any fixed set of parameters. We thus averaged the mating success of each tactic over 100 independent runs for each combination of these three variables: proportion of bafflers, population density, and habitat type. To check for the effect of the proportion of bafflers and population density on the mating success of baffling, we performed a linear regression analysis, with the mating success of bafflers as the response variable and the proportion of bafflers, population density and (proportion of bafflers)*(population density) set as fixed factors. The linear model was employed separately for the homogeneous landscape as well as the structured landscapes with different bush densities. To compare the mating success of the 3 ARTs, we constructed a linear model with mating success as the response variable, and the ARTs and proportion of bafflers in the population as fixed factors. The linear model was applied separately to simulation results from each combination of population density and habitat type. We then compared the intercepts for each of the ARTs to check for differences in mating success of the ARTs. Regression analyses and plotting were done in R 4.1.1 (R core team, 2022). Plotting used the ggplot2 (Wickham, 2016) package.

We clarify that the purpose of the statistical analyses is not to check the statistical significance as done in an empirical study; this is because by increasing the number of replications within our simulations, we can obtain an arbitrarily high degree of statistical significance (e.g., measured via p-values). Instead, our focus here is to quantify the relative strengths of mating success as a function of conditions such as population density and habitat structure.

### Sensitivity analysis

We varied some of the parameters to check if our model is sensitive to these parameters. The results of the sensitivity analysis are presented as supplementary information (Figures S6 to S15).

## Results

In the homogeneous habitat, the average mating success of baffling decreased with increasing proportion of bafflers in the population, implying negative frequency-dependent fitness benefits of baffling (Fig. 2A and 2C). The negative frequency dependence of baffling is strengthened as population density increases as indicated by the negative interaction of population density with the proportion of bafflers in the population (Fig 2D). In the homogeneous habitat, despite the strong negative frequency-dependent fitness effects on baffling, the average mating success of the baffling tactic was greater than the other ARTs (silent and calling) for all values of population density and proportion of bafflers that we tested in the simulations (Fig. 2 A). Further, the disparity between the mating success of baffling and the other ARTs increased with population density (Fig. 3A, B).

**Fig. 2.**
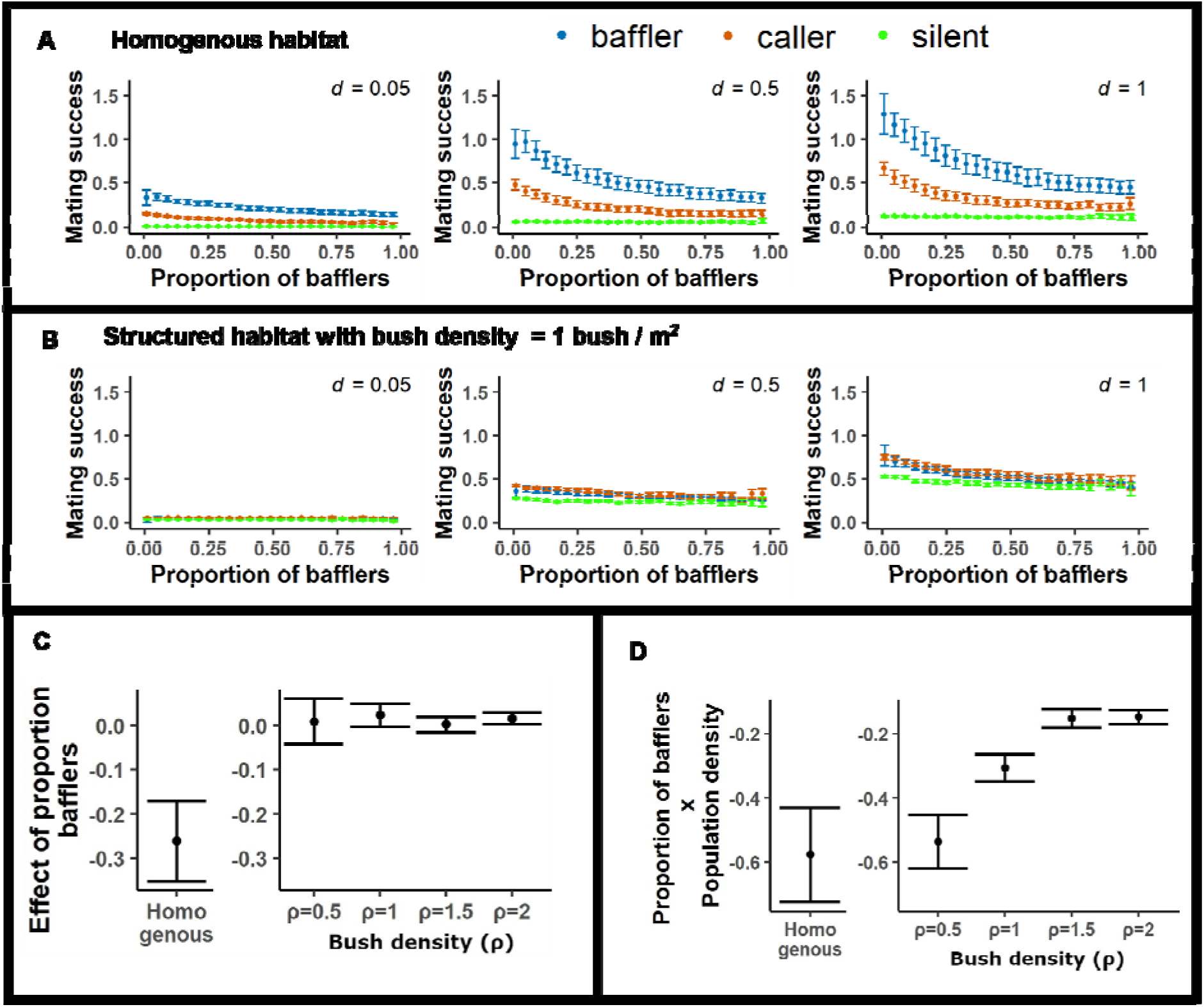
Mating success of ARTs is altered by spatial structure. A) Average mating success of the ARTs decreases with the proportion of bafflers in the homogenous landscape, with bafflers showing maximum success, for various population densities (*d*). B) Average mating success of the ARTs is altered dramatically in a structured landscape with bush density ρ=1 bush /m^2^, with nearly all strategies having similar success across baffler proportions and population densities. C) The strength of effect of proportion of bafflers on mating success of the baffling tactic across the homogeneous habitat and structured habitats having varying bush densities ρ. D) Interaction effects of proportion of bafflers and population density on the mating success of the baffling tactic across the homogeneous habitat and structured habitats having varying bush densities ρ. Error bars indicate 95% CIs. Figures A and B depict results for only a partial set of population densities and habitat type/bush densities (ρ). The complete results are presented in supplementary figure S1.

**Fig. 3.**
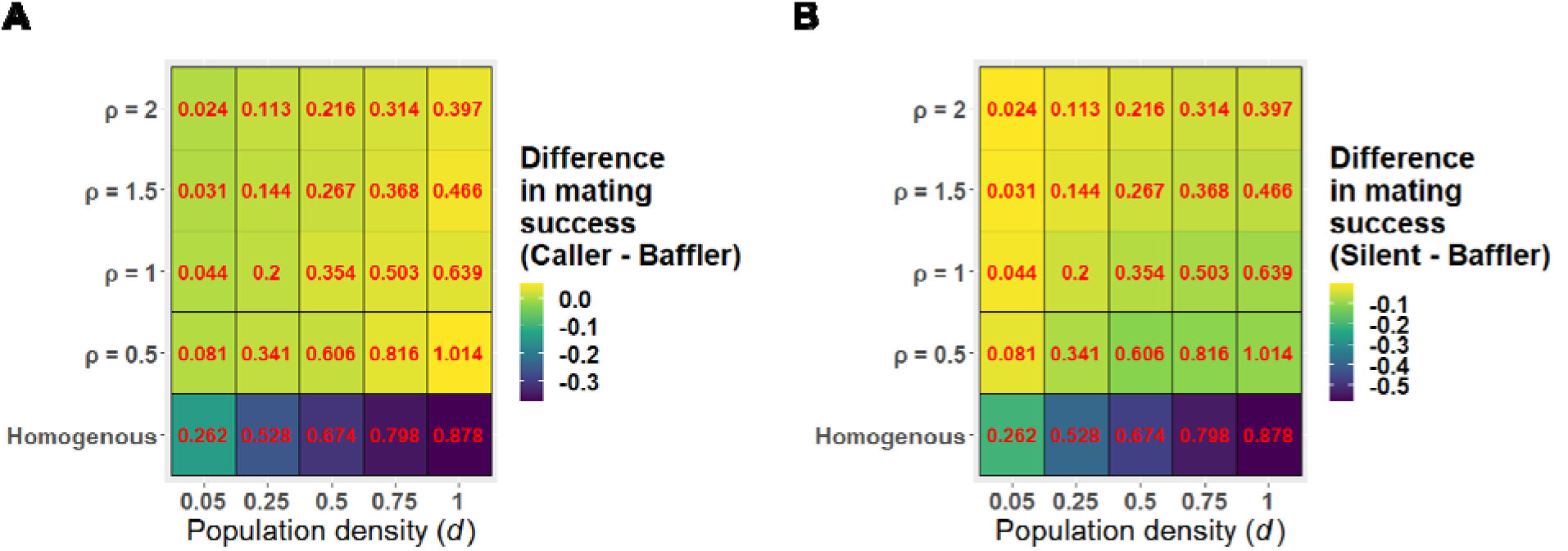
Disparity in mating success of tactics decreases in structured habitats compared to homogeneous habitats. A) Difference in mating success between calling and baffling tactic. B) Difference in mating success between silent and baffling tactic. Values (in red) represent mating success of baffling tactic to allow comparison of the difference in mating successes relative to the absolute mating success of baffling tactic.

When spatial structure in the form of bushes was introduced into the simulation landscape, we observed a weakening of the frequency dependence of the baffling tactic and the disparity between the mating success of baffling and the other tactics diminished considerably (compare Fig. 2A and 2B). The weakening of the strength of frequency dependence was independent of bush density in the structured habitat (Fig. 2B, 2C). However, increase in population density strengthened the frequency dependence of baffling at low bush densities (Fig. 2D).

While baffling was clearly superior in homogeneous habitats, in spatially structured habitats, the calling tactic showed marginally improved mating success compared to bafflers at all densities of bushes examined in the simulations (Fig. 3A). The difference between the silent and baffling tactics also reduced with the introduction of bushes. Increasing bush density and decreasing population density together decreased the disparity in mating success of the silent and baffling tactics (Fig. 3B). Sensitivity analyses revealed that the results remained qualitatively unchanged for various movement related (Fig. S6-S8; S10-S14) and demographic (Fig. S10 and S15) parameter values.

## Discussion

Demographic factors can affect the fitness outcomes of alternative reproductive tactics (ARTs). In the present study, we examined the effect of the proportion of bafflers, population density and habitat structure on the mating success of baffling males in the presence of other ARTs (calling and silent) expressed by males of a tree cricket by employing individual-based models (IbMs).

### Does baffling show negative-frequency dependence?

In homogeneous habitats, baffling showed clear negative frequency-dependence in mating success, and the strength of negative frequency-dependence increased with an increase in population density (Fig 2A). The negative frequency-dependence of mating success of bafflers can be attributed to increased competition for mates among bafflers as their frequency in the population increased. This is because, increasing the number of baffling males will increase competition for a fixed number of females, resulting in lowered per-capita mating success of bafflers. The increased strength of frequency-dependence with an increase in population density can be understood as an increase in the potential for competition: when population density is high, females are more likely to hear two or more male calls simultaneously and execute choice in phonotaxis. These results are similar to the negative frequency-dependence of mating success of calling males observed in caller-satellite systems (Rotenberry & Zuk, 2016; Rowell & Cade, 1993).

Interestingly, the introduction of spatial structure ameliorated negative frequency-dependence of baffling and led to more uniform mating success. This can be understood through the following reasoning: since movement across bushes is rare, most phonotaxis in our model happens within the same bush. The spatial scale over which bafflers directly compete for females is thus constrained by bush size, whereas in the homogeneous case, it is constrained by the size of the overlapping active sound fields of the calling males (*i.e.,* the distance up to which their call transmits before the amplitude falls below the behavioral threshold required for female phonotaxis). Thus, in our simulations, baffling males effectively compete mostly with other baffling males that co-occur on the same bush, whereas in the homogeneous cases, they compete with a larger number of baffling males that are present within their active acoustic spaces.

### Is baffling the superior tactic in structured habitats?

In the homogeneous case, despite the occurrence of negative frequency-dependence, the mating success of the baffling tactic was considerably higher than those of the other ARTs for all values of baffling proportion and population density we examined in our simulations (Fig 2A), in line with a previous simulation study (Deb et al., 2020). This is because bafflers enjoy a large increase in their call amplitude, which confers a twofold benefit to mating success by both increasing the distance from which bafflers can be detected and increasing their attractiveness when competing with other calling males that are close by. These effects together lead to bafflers having a much higher mating success than callers and silent males.

Introducing spatial structure decreased the absolute mating success of all three tactics compared to the homogeneous case (compare Fig. 2A and Fig. 2B). This can once again be understood by recognizing that bushes are small and across-bush flights are rare. Since across-bush movement is rare, most mating events are the result of phonotaxis that occurs towards a male in the same bush as the female. The advantages of baffling are limited in this case, since bafflers can primarily attract females that co-occur on the same bush. Given the low expected number of animals on a bush in structured habitats, the chance of a baffler co-occurring with a female on a bush is low, which limits the success of bafflers. Further, since females always move towards a sound that is sufficiently loud (in our simulations, > 60 dB SPL), the greater amplitude of bafflers is only advantageous in scenarios in which females are choosing between two or more alternative sources of sound, a scenario that is rare unless the global population density is high (Deb & Balakrishnan, 2014) and the habitat is unstructured.

### How does movement affect mate-encounter rate in structured versus homogenous habitats?

Individuals using tactics involving external structures such as leaves (in tree crickets) and burrows (in mole crickets) to increase the amplitude of their calls (Brandt et al., 2022) are often rendered immobile because these external structures are immovable. In our simulations too, bafflers were constrained to be stationary for the duration of each simulation run. In contrast, non-baffling callers, as well as silent males, moved around the landscape albeit with low likelihood. To test whether movement conferred any advantage to calling and silent males in homogeneous landscapes, we compared the results of our simulations where callers and silent males were allowed to move to those where they were not. The comparisons showed that movement did not provide any effective advantage to callers and silent males (Supplementary figure S2). Our results are in agreement with simulations investigating optimal mate search patterns in homogeneous habitats for mating systems that involve sexual signaling mates which show that signallers should move more slowly or less diffusively than the receivers to maximise their mate-encounter rate (Mizumoto & Dobata, 2018).

However, similar comparisons using structured habitats showed that callers and silent males showed higher mating success when they were allowed to move compared to when they were not allowed to move (Supplementary figure S3). Advantage due to movement may explain why in spatially structured habitats, calling males, despite their lower call amplitude showed slightly higher mating success than stationary bafflers (Fig. 2B, middle and right panel). Furthermore, the mating success of silent males in structured habitats (Fig. 2B) also increased compared to the negligible mating success they enjoyed in homogeneous habitats (Fig. 2A). Simulations with only silent males and females showed that mating success was higher in structured versus the homogeneous habitat, suggesting that spatial structuring increases mate encounter rate (Supplementary fig. S4). The higher random mate-encounter rate of silent males in structured habitats partially explains the reduction in the difference between the mating success of bafflers and silent males in structured habitats compared to the homogeneous habitat. Random mate encounter may have increased relevance to mating success in spatially structured habitats as females showed increased tendency to perform random movement relative to phonotaxis in structured habitats (Supplementary figure S5). The findings suggest that movement may facilitate mate-finding for both signaling and non-signaling males in structured landscapes, highlighting the importance of incorporating movement and spatial structure in future studies aiming to infer optimal mate-finding strategies in signal-receiver systems.

### The importance of habitat spatial structure for O. henryi

A recent study investigating the acoustic efficiency of different calling strategies in 3D landscapes using finite element analysis showed that baffling is more efficient in increasing the volume of the sound field than unaided calling for male *Oecanthus* species, which mainly call from vegetation (Brandt, et al., 2022). Despite the increased acoustic efficiency of baffling, however, only around 11% of *O. henryi* males baffle on a given night in natural field conditions (Deb, 2017). Deb et al. (2020) showed that baffling may be condition-dependent with smaller, lower intensity calling males opting to baffle as they stand to obtain a larger increase in mating success from baffling as compared to larger, louder males (Deb et al., 2020). Our results propose an additional explanation for the low percentage of bafflers in the field: calling may be more effective than baffling in structured landscapes as baffling highly constrains male movement. Additionally, it is evident from Fig. 3A that calling is more advantageous than baffling at high population densities with the reverse being true at low population densities across a wide range of bush densities. Thus, fluctuating population densities, which can be expected in a natural habitat, may allow the co-existence of the ARTs. Incorporating more ecological realism, such as saturating benefits (Deb et al., 2020) and energetic costs associated with loud signaling may further equalise the pay-offs of these ARTs, potentially explaining their coexistence. Predation risk, another relevant ecological factor, was found to be to too low to have meaningful effects on the coexistence of ARTs in *O. henryi* (Torsekar et al., 2019).

### Conclusion and future directions

Our study shows that incorporating spatial structure can affect the success of ARTs in a population. Firstly, while our finding corroborates previous modeling studies which demonstrate negative-frequency dependence of ARTs in crickets in unstructured landscapes, our study goesfurther to show that the negative frequency dependence breaks down in structured landscapes. Secondly, incorporating spatial structure reduces the disparity in mating success between ARTs. Post facto analysis of our findings revealed that movement played a pivotal role in reducing the disparity and the same is only observable in structured landscapes. Furthermore, in our model, baffling was more advantageous than calling in structured habitats and at low population densities. Thus, our study shows that population density plays a role in altering the hierarchy in mating success of ARTs in structured habitats. Summarily, by employing individual based modelling which allowed us to incorporate heterogeneity in the habitat as well as among individuals in a populations, we provide a first proof of concept for the role of spatial structure and movement ecology in facilitating the coexistence of signaling ARTs.

Analytic studies do propose cyclical dominance of strategies as a mechanism for the coexistence of 3 or more alternative strategies in nature (Szolnoki et al., 2014) with empirical support stemming from studies on biological systems ranging from side-blotched lizards (Sinervo & Lively, 1996) to bacteria (Kirkup & Riley, 2004). However, empirical insights suggest that cyclic dominance is quite rare in nature (Park et al., 2020), meaning the generality of cyclic dominance as a mechanism for coexistence is limited. Analytic studies of ecological coexistence in non-spatial models also suggest that the coexistence of multiple strategies is unstable (Alonzo & Calsbeek, 2010), unlikely (Han et al., 2012) or require special circumstances (Roy & Chattopadhyay, 2007). On the other hand, our study finds support in numerous other studies which show that habitat structuring can sustain the coexistence of competing species/phenotypes by promoting diversifying selection (Gray et al., 2008), demographic stochasticity (Calsbeek et al., 2002) or limiting interaction between competing individuals (Ursell, 2021).

We employed individual-based modelling in our study as it is a useful tool to model systems where relative importance of different factors and the nature of their interaction in determining the system dynamics is a priori unknown. IbMs can serve as a guide to building more sophistcated analytical models. For example, it would be interesting to see which factors of our model (such as caller:silent ratio, behavioural differences in mated versus virgin females, etc.) can be safely neglected in analytical approximations without affecting the qualitative behaviour of the system. Such analytical models, with their reduced complexity, may allow us to model long term evolutionary dynamics and thus predict persistence of ARTs

Although our study does not allow for trait evolution or demonstrate the coexistence of competing phenotypes over evolutionary time, it does show that incorporating habitat structure contributes to equalizing the mating success of competing ARTs. T There is a need for future studies to perform simulations over evolutionary timescales to examine the persistence or lack thereof of ARTs. It isimperative that these studies should consider spatial structure along with other important factors such as individual lifespan under natural conditions and energetic costs of signaling among others while examining the persistence of ARTs. Furthermore, mesocosm experiments that simulate homogenous and structured environments can be employed to test the efficacy of signal-based ARTs in obtaining mates. Individual behavioural tracking in these mesocosm experiments can provide longitudinal data on the per-capita expression rates of the ARTs along with their corresponding mating successes. Comparing the mating success of ARTs in both homogenous and heterogenous mesocosms will help test the hypothesis whether habitat structuring indeed contributes to the reduction in disparity of mating success associated with the ARTs. Furthermore, experiments that incorporate increasing levels of heterogeneity can be used to test whether the stabilizing effect of habitat structure on the fitness of ARTs increases with habitat heterogeneity. Such studies can inform conservation decisions concerned with the modification of landscapes housing organisms that display ARTs.

## Acknowledgements

We acknowledge infrastructure support from DST-FIST and DBT-IISc partnership program, Government of India. MAS is supported by the Prime Minister’s Research Fellowship from the Ministry of Education, Government of India. ASB was supported by a Kishore Vaigyanik Protsahan Yojana (KVPY) fellowship (ID: SX-1711025) from the Department of Science and Technology, Government of India. We thank the anonymous reviewers for their insightful comments that have helped improve the manuscript.

## Author Contributions

**MAS:** Conceptualization, Methodology, Software, Validation, Formal analysis, Investigation, Data Curation, Writing – Original Draft, Writing – Review & Editing, Visualization; **ASB:** Conceptualization, Methodology, Software, Validation, Formal analysis, Investigation, Data Curation, Writing – Original Draft, Writing – Review & Editing, Visualization; **RB:** Conceptualization, Methodology, Writing – Review & Editing, Supervision; **VG:** Conceptualization, Methodology, Writing – Review & Editing, Supervision

## Open Research statement

All data and codes generated in the production of this manuscript can be found on a public GitHub repository at: https://github.com/ThePandalorian/Sadiq_et_al_2023_oecanthus_ART_IbM

